# *Streptococcus pneumoniae* accessory capsular genes modulate fitness, pathogenicity and immune evasion

**DOI:** 10.1101/2025.09.11.675585

**Authors:** Jia Mun Chan, Helen Flynn, Suji Kim, Hayley Demetriou, Elisa Ramos-Sevillano, Amal Alsari, Lucy Roalfe, Giuseppe Ercoli, Akuzike Kalizang’oma, Comfort Brown, Chrispin Chaguza, Neil French, Jeremy Brown, David Goldblatt, Luiz Pedro Sorio de Carvalho, Robert S. Heyderman

## Abstract

Globally, *Streptococcus pneumoniae* disproportionally affects children in resource-poor settings, older adults and people living with HIV. Frequently found as an asymptomatic colonizer of the nasopharynx, this versatile pathogen is a prominent cause of pneumonia, meningitis, bacteraemia and otitis media. Recently, a serotype 3 capsule variant (GPSC10-ST700) has expanded in Malawi with enhanced vaccine escape potential. Here, using a mutational and complementation approach, we show that loss of accessory capsular genes in GPSC10-ST700 contribute to increased opsonophagocytic resistance in this lineage. Although originally thought to be nonfunctional pseudogenes, we show that these genes modulate fitness and the global phosphoproteome in serotype 3 strains. These findings highlight that vaccine escape may be mediated through variations in the pneumococcal capsular locus that enhance fitness, pathogenicity and immune evasion, without capsule switching.

**IMPORTANCE:** Pneumococcal polysaccharide-conjugate vaccines (PCV) target the polysaccharide capsule (CPS), which is a dominant virulence factor. However, current PCVs induce suboptimal protection against serotype 3 strains, which produce a thicker capsule that when released from the bacterial surface, interferes with antibody-mediated bacterial killing and protection. We recently described the clonal expansion of a sequence type (ST) 700–GPSC10 serotype 3 lineage in Malawi post-PCV13 introduction. This lineage is characterized by the absence of at least 6 genes in its *cps* locus and a distinct antimicrobial resistance (AMR) profile compared to other serotype 3 strains. Here we uncovered a functional role for the accessory capsular genes (*acl*) in serotype 3, previously considered to be pseudogenes, which modulate capsule production, shedding, serum tolerance, and bacterial fitness. By linking genotype to phenotype, our work provides new insights into the molecular basis of serotype 3 immune evasion, informing the design of more effective pneumococcal vaccines.

## INTRODUCTION

*Streptococcus pneumoniae* is a highly adaptable pathobiont. Pneumococcal infections are responsible for the death of approximately 300,000 children under the age of 5 each year (1). The pneumococcal polysaccharide capsule (CPS) is a key virulence factor that protects the bacterium from desiccation, antibody- and complement-mediated immune clearance, and it serves as an important carbon source for the bacteria during periods of starvation (2–9). Indeed, unencapsulated strains of *S. pneumoniae* only rarely cause invasive disease (10, 11). Pneumococcal conjugate vaccines (PCV) targeting the capsule polysaccharide have been introduced into the routine infant immunisation programme in at least 155 countries, successfully reducing the burden of disease (12–14).

The pneumococcus has over a hundred biochemically and antigenically distinct capsule serotypes; however, 20–30 serotypes are responsible for most of the invasive pneumococcal disease (IPD) (15, 16). All but two serotypes use a *Wzy*-dependent pathway to produce and anchor their capsule to the peptidoglycan layer (17–19). The capsular polysaccharides of serotype 3 and serotype 37 strains are made through a synthase pathway and are loosely anchored to the cell by a covalent bond with phosphatidylglycerol (20). This loose capsule adherence gives serotype 3 strains their distinctive mucoid phenotype when plated on blood agar plates (Figure 1).

**Figure 1.**
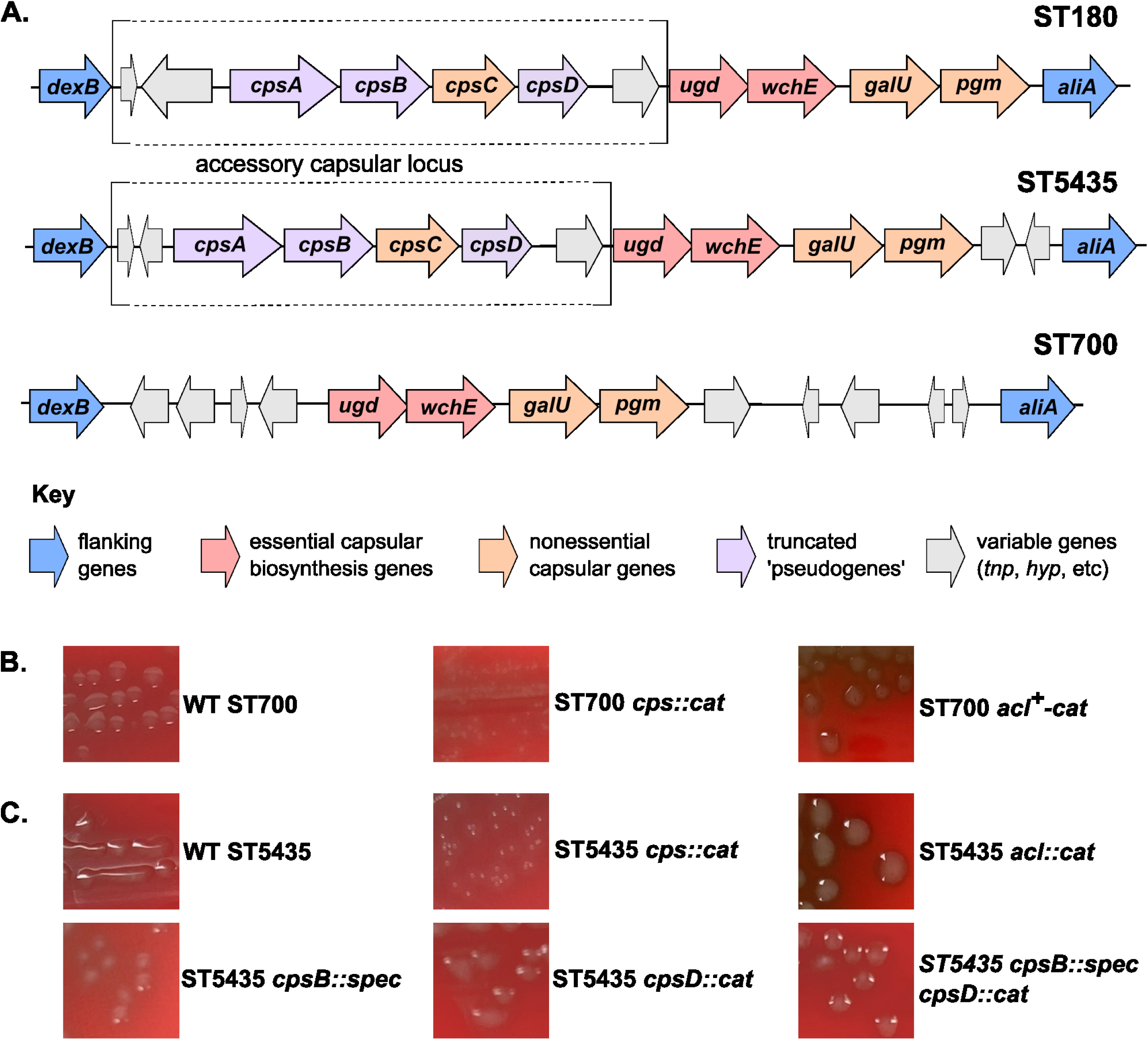
Serotype 3 ST700 strains lack multiple accessory capsular genes, with no overt effect on capsule production. (A) Genetic organization of the capsular locus in three serotype 3 lineages: ST180 (reference genome (24)), ST5435, and ST700; not to scale. The variable genes immediately downstream of *dexB* and upstream of *ugd* are conserved in ST180 and ST5435. (B) Allelic replacement of all genes between *dexB* and *aliA* in ST700 (*cps::cat*) abolishes mucoid (capsule) production, while reintroduction of the accessory capsular locus (*acl*) genes does not. (C) Deletion of *acl* genes does not impact mucoid (capsule) production in ST5435.

Real-world vaccine effectiveness against serotype 3 has been lower than predicted through clinical trials (12, 13, 21). This has been attributed to lower amounts and lower functional effectiveness of vaccine-elicited antibodies than expected, partly attributed to the ability of serotype 3 strains to release capsule into the extracellular milieu (21–23). In a mouse model of pneumococcal septicaemia, shed serotype 3 capsule blocks the protective effect of passive immunization (22) Flanked by the *dexB* and *aliA* genes, the *S. pneumoniae* capsular biosynthesis locus is hypervariable (24, 25). We and others have identified at least three variations of the serotype 3 CPS locus (Figure 1A) (24, 26, 27). The shortest version, which is encoded by an emergent GPSC10-ST700 lineage in Malawi, lacks the first six genes of the locus, collectively termed accessory capsular locus (*acl*) (Figure 1) (27, 28). Two of these missing genes are truncated homologs of *cpsB* (*wzh*) and *cpsD* (*wze*), which are part of a bacterial tyrosine kinase (BYK) system that regulates capsule expression and chain length (29–32). Where these truncated *cpsB* and *cpsD* homologues have been present in serotype 3 strains, they have been considered to be pseudogenes and therefore nonfunctional (33, 34). However, we have shown that GPSC10-ST700 isolates are more resistant to opsonophagocytic killing (OPK) than the previously dominant circulating serotype 3 lineage in Malawi, GPSC9-ST5435 (27). This has led us to speculate that the *cpsB* and *cpsD* homologues are functional, and that their loss from the GPSC10-ST70 *cps* locus results in enhanced OPK resistance and virulence through altered enzyme phosphorylation and capsule shedding.

Here we show that *acl* genes, in particular *cpsB,* influence OPK susceptibility in serotype 3 through the regulation of capsule production and shedding. Mutating *cpsB* and *cpsD* shifts the global phosphoproteomic landscape and slows *S. pneumoniae* growth*, c*ollectively confirming that these *acl* genes are not pseudogenes but modulate both fitness and immune evasion in *S. pneumoniae* serotype 3.

## RESULTS

### Deletion or reintroduction of *acl* does not alter colony morphology

We deleted all six *acl* genes from the serotype 3 GPSC9-ST5435 strain BVY11Z (hereafter WT ST5435) and reintroduced the genes into the capsular locus of serotype 3 GPSC10-ST700 strain BVY23H (hereafter WT ST700) (Figure 1A (27), using an allelic replacement strategy (detailed in Materials and Methods). In concordance with previous studies (33, 34), we did not see any obvious differences in colony morphology between the WT ST5435 and its *acl* deletion mutant or between WT ST700 and its *acl+* reintroduction mutant (Figure 1B-C)(33, 34). By contrast, deletion of the entire capsular locus between the conserved flanking genes *dexB* and *aliA* in both ST5435 and ST700 resulted in the loss of the mucoid phenotype characteristic of encapsulated serotype 3 strains (Figure 1B-C).

Within the *S. pneumoniae* serotype 3 capsular locus, *cpsB* and *cpsD* have been thought to be nonfunctional pseudogenes as they are (i) truncated compared to their homologues in other serotypes, (ii) dispensable for capsule production, and (iii) not to be expressed during growth (18, 33, 35). Indeed, deletion of *cpsB* and *cpsD*, either singly or in tandem, did not alter colony morphology by the serotype 3 strain (Figure 1C). The characteristic serotype 3 mucoid phenotype is generally considered to be evidence of capsule production and therefore, as expected, *cpsB* and *cpsD* are not essential for capsule production in serotype 3. Serotyping of GPSC10-ST700 isolates using latex agglutination also showed positive identification as serotype 3 strains (28), and as such *cpsB* and *cpsD* do not alter the immunodominant epitope of serotype 3 capsule.

### Changes in the *acl* region of Serotype 3 strains influence opsonophagocytic resistance

To evaluate the role of the serotype 3 *acl* in modulating susceptibility to opsonophagocytic killing (OPK), we subjected WT ST700 and its *acl* reintroduction mutant (ST700 *acl*^+^-cat), as well as WT5435 and its *acl* deletion mutant (ST5435 *acl::cat*) to a WHO standard OPK assay using pooled serum from PCV-immunised adults (36, 37). In concordance with our prior study, WT ST700 was more resistant to OPK than WT5435 (Table 1) (27). However, contrary to our expectations, not just the ST700 *acl+* but also the ST5435 *acl*-strains were more susceptible to killing than their respective WT strains (Table 1). To determine if the BYK capsule regulation system plays a role in OPK susceptibility, we additionally tested the ST5435 *cpsB* deletion mutant (ST5435 *cpsB::spec*) and ST5435 *cpsD* deletion mutant (ST5435 *cpsD::cat*) in the OPK assay. Deletion of *cpsB*, which encodes a putative phosphatase, but not *cpsD*, which encodes a putative autophosphorylating kinase, greatly increased susceptibility of ST5435 to OPK killing (Table 1). To determine if this phenotype is conferred by increased complement deposition in the ST5435 *cpsB* and *acl-* mutants, we used flow cytometry to measure C3b deposition. However, we did not observe significant differences in the amount of C3b deposition between the mutants and the WT strains (Supplementary Figure 1).

**Table 1.** Loss of *acl* genes does not increase opsonophagocytic killing (OPK) resistance in Serotype 3 strains. The assay was performed using a WHO-standard operating protocol in an approved laboratory, using pooled serum from PCV-immunized individuals. A higher opsonophagocytosis titre value indicates increased susceptibility to OPK (more killing compared to a reference strain).

| Genotype | Opsonophagocytosis Index |  |
| --- | --- | --- |
|  | *Experiment 1* | *Experiment 2* |
| WT ST700 | 50 | 47 |
| ST700 *acI*+ | 123 | 166 |
| WT ST5435 | 99 | 76 |
| ST5435 *acI::cat* | 146 | 196 |
| ST5435 *cpsB::spec* | 191 | 165 |
| ST5435 *cpsD::cat* | 90 | 111 |

### Changes in the *acl* region affect the amount of capsule produced and released

Unlike most other pneumococcal serotypes, the serotype 3 capsule is not attached to the peptidoglycan but is instead covalently linked to phosphatidylglycerol (20). Shed serotype 3 capsule protects the bacteria from killing in both laboratory OPK assays and in a murine infection model (22). We therefore hypothesized that WT ST700 would shed more capsule compared to WT ST5435. We measured the amount of capsule expressed and shed by WT ST700 and its *acl+* reintroduction mutant, WT ST5435 and its *acl-* deletion mutant, its *cpsB, cpsD* and *cpsB cpsD* double mutant. The unencapsulated mutants of both ST700 and ST5435 were used as negative controls.

When grown in rich liquid medium (BHI), the two WT strains shed similar amounts of capsule, with notable variations (Figure 2A). The ST5435 *acl-* and *cpsB-* mutants, as well as the ST700 *acl+* reintroduction mutant, tended to shed lower amounts of capsule compared to both WT strains, but the differences were not statistically significant (Figure 2A). Furthermore, we found that WT ST5435, ST5435 *acl-, cpsB, cpsD, cpsB cpsD* double mutants, or the ST700 *acl+* reintroduction mutant shed most of their capsule when grown in BHI (Figure 2A-B).

**Figure 2.**
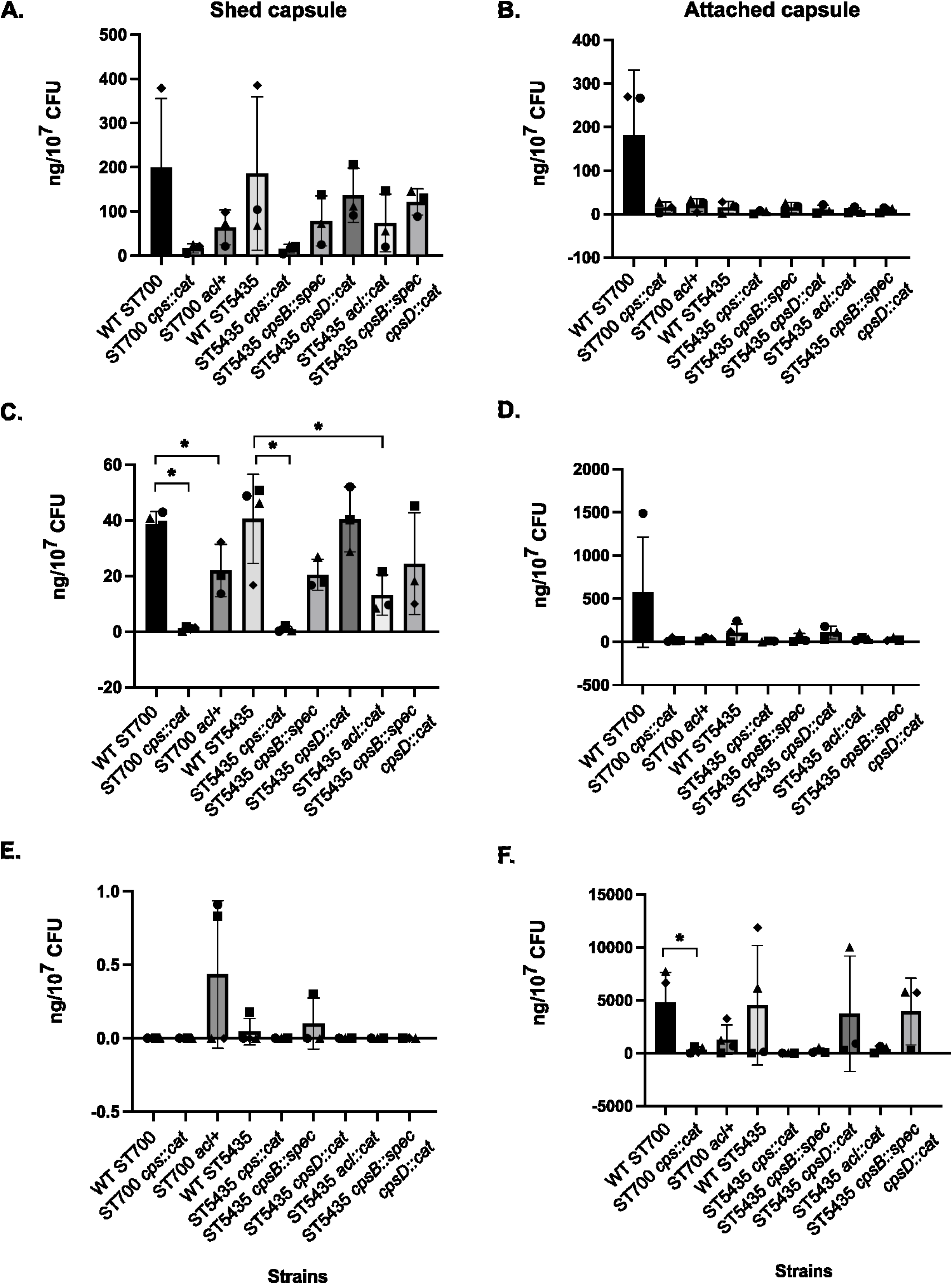
Perturbation to the *acl* region negatively impact Serotype 3 ability to react to normal human serum (NHS) treatment. The amount of shed capsule (A,C,E) and attached capsule (B,D,F) made by various Serotype 3 strains during growth in BHI to OD_600_ of 0.3 (A,B), without (C,D) and in the presence (E,F) of serum stress for an additional hour were quantified using ELISA and normalized to CFU counts at the time of harvest. Unencapsulated mutants (*cps::cat*), which do not produce capsule, were used as a control for off-target antibody binding. The serum stress experiment was performed at least three times on different days, and ELISA was performed with technical duplicates. Statistical comparisons were performed using Mann-Whitney test; * indicates p-value < 0.05.

To test whether the amounts of shed and attached capsule vary under immune pressure, we grew the strains for an additional hour with supplementation of either 20% normal human serum (NHS, purchased from Merck) or an equal volume of fresh BHI media without serum. The overall patterns of shed and attached capsule by the various strains were similar in the pre-treatment and the 0% NHS conditions (Figure 2A-D). In contrast, we detected little to no shed capsule from any of the strains grown in 20% NHS (Figure 2E), with significant increases in the amount of capsule retained on the cell surface by WT ST700, WT ST5435 and its *cpsD* and *cpsB cpsD* double mutant (Figure 2F). The ST700 *acl+* reintroduction mutant, ST5435 *acl-* and *cpsB* deletion mutant showed non-significant increases in attached capsule when grown in 20% NHS treatment (Figure 2F). *S. pneumoniae* is not susceptible to killing by complement components in the absence of phagocytes (38). All our strains, including the unencapsulated mutants continued to grow in the presence of 20% NHS (Supplementary Figure 2). Overall, although there were extensive day-to-day variations in capsule expression, the strains that were more susceptible to OPK killing showed lower levels of attached capsule when grown in 20% NHS (Table 1, Figure 2F). This suggests that the ability of ST700 to increase the amount of attached capsule in response to serum stress is negatively affected by the presence of the *acl* region, while the reverse is true for ST5435.

### Changes in the *acl* region affect pneumococcal growth

To determine if changes in the *acl* regions also affect pneumococcal growth, we conducted overnight growth assays using BHI as nutrient media. The reintroduction of *acl* into ST700 resulted in a small but reproducible half-hour decrease in the lag time compared to WT ST700 (Figure 3A). By contrast, deletion of *acl* in ST5435 resulted in a small but reproducible half-hour increase in lag time compared to WT ST5435 (Figure 3B). Mutation of *cpsB*, *cpsD,* or both together also increased the lag time compared to WT ST5435 (Figure 3B). Deletion of the entire *cps* region led to a small but reproducible decrease in lag time for WT ST700 but not WT ST5435, with a lower maximum OD_600_ for the unencapsulated mutants compared to their WT strains (Figure 3C-D). Overall, the presence of *acl* promoted the growth of both ST700 and ST5435 strains in laboratory media.

**Figure 3.**
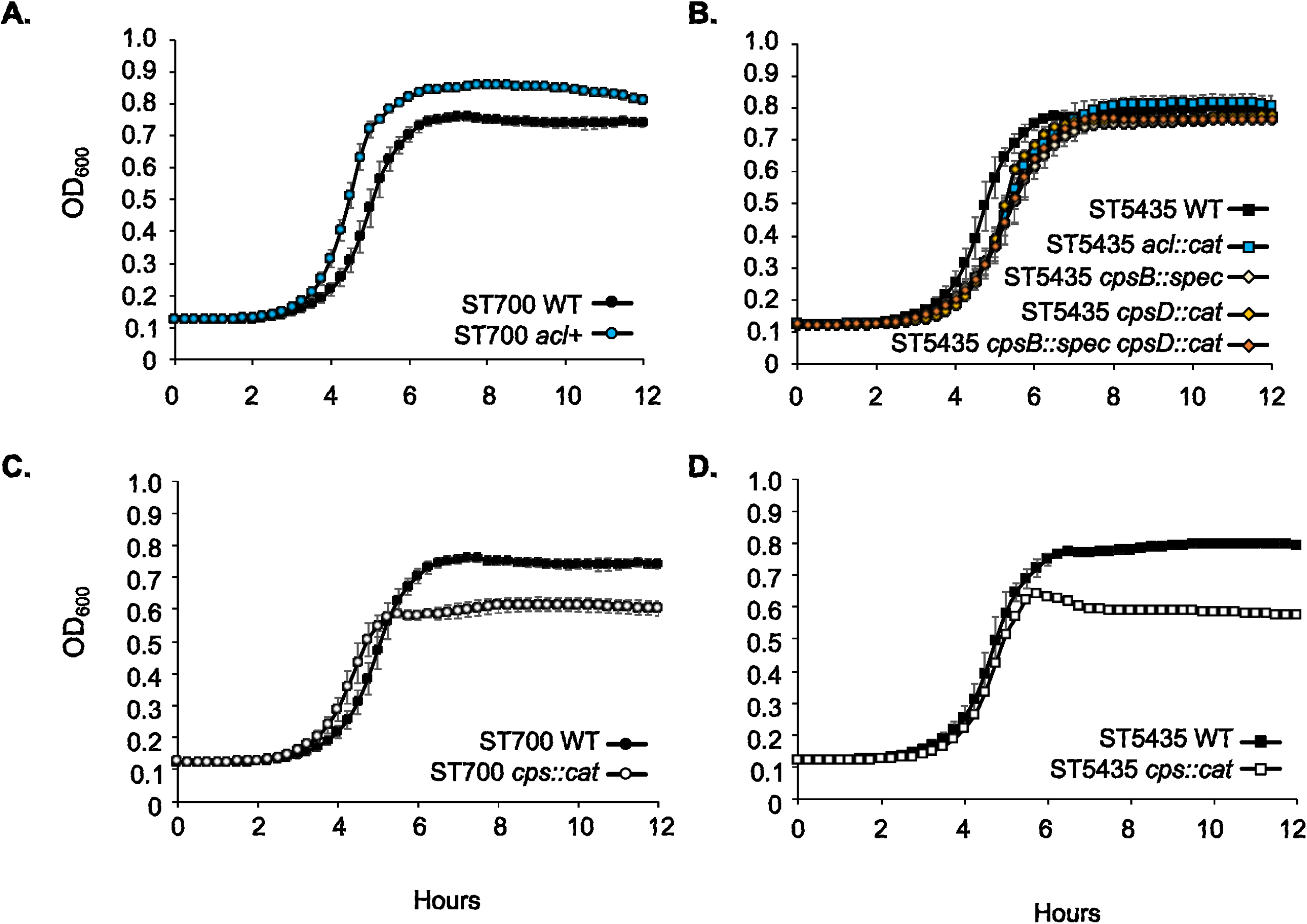
Disruption of the capsular locus alters growth kinetics. A) Reintroduction of *acl* into ST700 decreases the lag time and increases the maximum OD of the strain. B) Mutation of *cpsB, cpsD,* both genes together or the entire *acl* region in ST5435 increases the lag time of the bacterium. C) Mutation of the entire *cps* locus in ST700 decreases the lag time in a small but reproducible manner and decreases the max OD of the strain. D) Mutation of the entire *cps* locus in ST5435 reduces the max OD of the strain without affecting lag time. The experiment was performed on three separate days with technical triplicates (n=3); graphs show the averaged OD_600_ reading with standard deviation as error bars.

### Presence of BYK-encoding genes alters the pneumococcal global phosphoproteome

Since *cpsB* and *cpsD* encode putative BYK components, we performed phosphoproteomic analysis to identify possible BYK system targets in serotype 3 strains. WT ST700 and its *acl+* reintroduction mutant, WT ST5435 and its single *cpsB, cpsD,* and double *cpsB cpsD* mutants were grown to early log phase, treated with NHS or equal volume fresh medium and subsequently harvested for phosphoproteomics processing and LC-MS/MS analyses. The obtained mass spectra were searched against a custom database comprising predicted proteins from the whole-genome sequences of WT ST700 and WT ST5435 strains. Proteins may have more than one phosphorylation site, and each site may be phosphorylated independently. This results in multiple phosphopeptides mapped to the same protein present on both sides of the volcano plot (Figure 4, Supplementary Table 1; see Tuf as an example).

**Figure 4.**
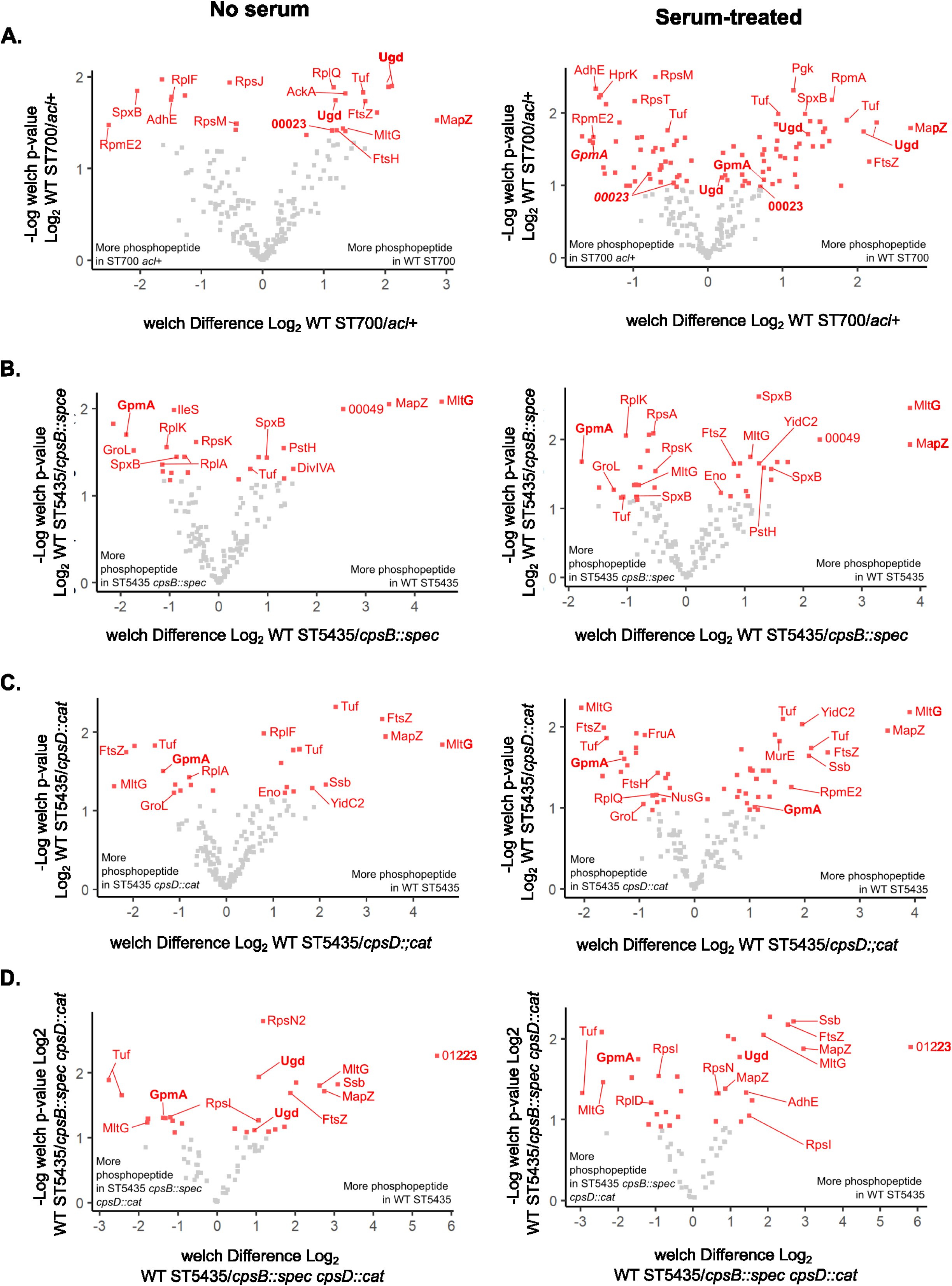
Changes to the *acl* region impacts global phosphoproteome patterns. WT ST700 and its *acl* reintroduction mutant (A), and WT ST5435 and its *cpsB* (B), *cpsD* (C), and *cpsB cpsD* (D) double mutants were grown in rich media with and without 20% normal human serum for an hour before harvesting for phosphoproteomic analysis. Phosphopeptides were enriched using Sequential Metal Oxide Affinity Chromatography strategy and subjected to LC-MS/MS. Each square represents a unique phosphopeptide. Since a protein may have more than one phosphorylation site, multiple phosphopeptides may be mapped to the same protein. Red squares indicate unique phosphopeptides that were significantly enriched in one strain compared to the other. **Bold** indicate phosphorylation of proteins of interest (Ugd, GpmA, BNFDFNGKA_00023 (annotated only as 00023)) at a serine or threonine residue. ***Bold italics*** indicate phosphorylation at a tyrosine residue.

First, we analysed the two Serotype 3 strain-essential capsular biosynthesis genes within the locus, *ugd* and *wchE* (Figure 1A). Ugd is a UDP-glucose dehydrogenase and WchE is the capsular polymerase (17). We detected higher levels of Ugd phosphorylation in WT ST700 compared to its *acl+* mutant in both serum-treated and serum-free growth conditions (Figure 4A, Supplementary Table 1; annotated in genome as *ugd_3*). However, the phosphorylation sites detected are not tyrosine but were instead serine and threonine (Supplementary Table 1). We also detected differential Ugd phosphorylation on a threonine residue in the WT ST5435-*cpsB cpsD* double mutant pair but not with the WT ST5435-*cpsB* or WT ST5435-*cpsD* single mutant pairs (higher in WT; Figure 4B-D, Supplementary Table 1; annotated in genome as *ugd_1*). We did not detect differential phosphorylation of WchE, or any other proteins encoded within the capsular locus, including *acl* gene products.

Reintroduction of *acl* into ST700 led to significant shifts in the global phosphoproteome, especially in response to serum treatment (Figure 4A). There were higher levels of phosphorylation of cell division-related proteins (MltG, FtsZ) in WT ST700 and higher levels of phosphorylation of multiple ribosomal proteins in the ST700 *acl+* reintroduction mutant (Figure 4A). A similar pattern with cell division-related proteins was seen when comparing the phosphoproteome of WT ST545 and its mutants (Figure 4B-D). Mutation of *cpsD* led to larger shifts in the phosphoproteome compared to mutation of *cpsB* under serum stress conditions (Figure 4B-C, Supplementary Table 1).

We identified two proteins that were differentially phosphorylated on a tyrosine residue, GpmA and hypothetical protein BNFDFNGKA_00023, which were enriched in the ST700 *acl+* mutant under serum stress (Figure 4A, Supplementary Table 1). GpmA is also phosphorylated at multiple serine and threonine residues, and we detected higher amounts of a GpmA serine phosphopeptide in WT ST700 compared to its isogenic *acl*+ mutant, Conversely, we detected higher amounts of the GpmA serine phosphopeptide in all the ST5435 mutants compared to WT ST5435 in all growth conditions (Figure 4, Supplementary Table 1). GpmA is a 2,3-bisphosphoglycerate-dependent phosphoglycerate mutase involved in glycolysis, with pleiotrophic effects in other cellular function (39, 40). While CPS can be used as a carbon source in nutrient-limited conditions, GpmA’s role in CPS catabolism or biosynthesis is unknown (2).

BNFDFNGKA_00023 is a hypothetical protein with homology to cell filamentation protein Fic and contains a FIDO protein-threonine AMPylation domain (41). Phosphorylation of target proteins by FIDO domain proteins tend to disrupt the signalling or function of the protein, and Fic proteins tend to target GTPases (41). It is probable that the global shifts in the phosphoproteome seen with the ST700 *acl*+ mutant is partially facilitated by the function of BNFDNGKA_00023. We also detected BNFDNGKA_00023 phosphopeptide in the non-serum treated ST700-ST700 *acl+* samples; however, the difference was not statistically significant (Supplementary Table 1). We did not detect phosphorylation of the associated ST5435 homolog (KOFMOAC_00327) in our dataset (Supplementary Table 1).

Collectively, our results suggest that the *acl* genes exert an indirect and broad-ranging effect on the global phosphoproteome patterns. Given the preponderance of cell division and ribosomal proteins affected in our dataset, it is likely that these shifts contribute to the differential growth patterns in the mutants compared to WT strains.

## DISCUSSION

There is increasing evidence that the introduction of PCV13 has achieved little to no population-level immunity to serotype 3, resulting in ongoing carriage and considerable residual disease (12–15, 21, 42). Our previous identification of a GPSC10-ST700 serotype 3 lineage that has expanded in Malawi and South Africa provides the opportunity to better understand the link between variations in the serotype 3 *cps* locus and immune evasion (27, 28). This lineage is characterized by the absence of at least 6 *acl* genes near the start of the CPS locus and by increased resistance to opsonophagocytic killing (27).

In this study, we found that reintroducing the *acl* region into GPSC10-ST700 reduced resistance to opsonophagocytic killing. While this suggests that the absence of the *acl* confers a selective advantage under vaccine pressure, particularly where immunogenicity is incomplete, mutating *acl* or just *cpsB* alone in a serotype 3 ST5435 background (less intrinsically serum resistant than ST700) confers the same reduction in resistance to opsonophagocytic killing. Our phosphoproteomic analysis links GpmA and its role in glycolysis (intimately linked to pneumococcal capsule biosynthesis (43)) to *cpsB* and *cpsD* functionally; however, the molecular basis of resistance to OPK is complex, likely involving genes within and outside of the capsular biosynthesis locus. For example, cellular rather than capsular phosphoglucomutase (*pgm*) has been shown to play a more prominent role in serotype 3 capsule biosynthesis (34). Additionally, detection of BNFDNGKA_00023 as a differentially phosphorylated protein in the ST700 background but not in the ST5345 strains suggest that additional mutations distal to the capsule locus contribute to the serum resistance of ST700 compared to ST5435.

As previously reported with serotype 3 strain WU2 (22), all our encapsulated serotype strains tested were capable of shedding capsule into the supernatant, and production of capsule was upregulated in response to serum stress. Overall, WT ST700 produces and retains more capsule polysaccharide on its cell surface in the absence of serum stress compared to all the other serotype 3 strains tested, potentially enhancing resistance to OPK and survival during infection. In general, strains that were more susceptible to opsonophagocytic killing (ST700 *acl+,* ST5435 *acl-*, ST5435 *cpsB*) did not increase capsule production in response to serum stress in our experiments (Figure 2F, Table 1). It is noteworthy that there was substantial day-to-day variation in the amount of capsule shed or retained on the bacterial cell surface. While it is hypothesized that the loose anchoring of the serotype 3 capsule to the cell surface facilitates capsule release, the mechanism and regulation of capsule release remain unknown. *S. pneumoniae* is phase variable (25, 44–46), leading us to speculate that this heterogeneity may reflect a bet-hedging strategy.

Indeed, there has been previous reports that phase variation in capsule levels and colony opacity modulate biofilm formation, nasopharyngeal colonization and virulence (45, 47–50). Our data showing altered capsule release patterns in the presence of normal human serum, as well as the observation that the pneumococcus downregulates surface capsule expression to facilitate epithelial invasion (51, 52), further supports this possibility.

We also demonstrate a link between the *acl* and bacterial growth potential, with reintroduction of *acl* into ST700 resulting in a half-hour decrease in the lag time compared to WT ST700, and deletion of *acl* in ST5435 or mutation of *cpsB* or *cpsD* (previously thought to be nonfunctional pseudogenes) resulted in a half-hour increase in lag time compared to WT ST5435. While our phosphoproteomic analysis suggests that the *acl* genes largely exert an indirect effect on capsule status, the preponderance of cell division and ribosomal proteins in the global phosphoproteome suggests that their differential phosphorylation contributes to their differing growth patterns. We have previously shown that serotype 3 GPSC10-ST700 is tetracycline resistant, has a higher MIC for penicillin than other serotype 3 lineages isolated in Malawi, appears to have a propensity for capsule switching, and possesses virulence genes involved in pathogenicity and colonization (27). It is possible that changes in the global phosphoproteome driven by *acl* genes also contribute to the above phenotypes.

In serotype 2 strains, where most of the detailed biochemical studies of CPS synthesis and regulation have been done, CpsB dephosphorylates the autophosphorylating kinase CpsD under specific conditions to exert changes to CPS length (29–32). It is therefore reasonable to expect that CpsD phosphorylation would increase in the absence of *cpsB*. Surprisingly, we were unable to detect differential CpsD phosphorylation in our phosphoproteomic experiments. It is possible that CpsD does not function as a kinase, and CpsB does not act upon CpsD in serotype 3 strains. It is also possible that our approach is limited in its sensitivity, as many of our hits are associated with highly abundant proteins across pneumococcal strains. Likewise, we cannot discount the possibility that one or more *acl* genes are not translated, and that transcription of the genes itself exerts a regulatory effect. Indeed, there is ample evidence of sRNA regulating processes in *S. pneumoniae* (53).

Regardless, our data showed that alterations to the *acl* region and specific allelic replacement of *cpsB* and *cpsD* exert a global effect on the bacterial phosphoproteome. Future biochemical analysis of these gene products would help elucidate their mechanism of action.

The *acl* genes have long been considered nonfunctional pseudogenes in serotype 3 strains (17, 18). Together, our data challenge this assumption and demonstrate that *acl* genes function to modulate fitness, pathogenicity, and immune evasion in ST5435 and in ST700, at least in part by regulating capsule production and shedding. In the context of increasing reports of capsular locus variants (genetic changes that do not result in a serotype change) expanding in populations where PCVs have been introduced, understanding how these genetic changes might confer a competitive advantage is essential (54–58). The direct and indirect effects of mutations in the pneumococcal BYK system located in the *cps* are likely to have widespread effects on pathogenesis.

## MATERIALS AND METHODS

### Bacterial growth

*Streptococcus pneumoniae* strains were grown on Columbia agar base with 5% defibrinated horse blood (EO Labs) or statically in Brain-Heart-Infusion (BHI) broth (Oxoid) or Todd-Hewitt broth (Oxoid) supplemented with 0.5% yeast extract (THY) at 37°C, 5% CO_2_. For mutant selection, where appropriate, growth medium was supplemented with antibiotics at the following concentrations: chloramphenicol (10 µg/ml), spectinomycin (100 µg/ml). Working stocks were prepared by freezing THY cultures at an OD_600_ of ∼0.4 with 10% glycerol. The freezing medium was removed upon thawing, and bacterial cells were resuspended in the desired medium. A complete list of strains used in this study is summarized in Table 2.

**Table 2.** List of strains and plasmids used in this study.

| Designation | Genotype | Source |
| --- | --- | --- |
| *Strains* |  |  |
| WT ST700<br>(BVY11Z) | Serotype 3 ST700 clinical isolate | (27, 28) |
| WT ST5345<br>(BVY23H) | Serotype 3 ST5435 clinical isolate | (27, 28) |
| AA1 | ST700 *cps::cat* | This work |
| AA2 | ST5435 *cps::cat* | This work |
| ECSPN300 | ST5435 *cpsB::spec* | This work |
| ECSPN301 | ST5435 *acl::cat* | This work |
| ECSPN302 | ST5435 *cpsD::cat* | This work |
| ECSPN303 | ST5435 *cpsB::spec cpsD::cat* | This work |
| ECSPN306 | ST700 *acl<sup>+</sup>-cat* | This work |
| *Plasmids* |  |  |
| pEMcat | Minitransposon plasmid; source of *cat* (*cm<sup>R</sup>*) cassette | (60) |
| pR412 | Plasmid derived from ColE1; source of *spec* (*spec<sup>R</sup>*) cassette | (61) |

### Genetic manipulation and strain construction

Genetic manipulation of *S. pneumoniae* was carried out using a competence-stimulating peptide (CSP)-mediated transformation assay (59). Briefly, strains were grown in 12 mL of THY pH 6.8 supplemented with 1 mM CaCl_2_ and 0.2% BSA at 37°C with 5% CO_2_ to an OD_600_ of 0.01-0.03. Cultures were then harvested and resuspended in 1 mL of pre-warmed THY pH 8.0, supplemented with 1 mM CaCl_2_ and 0.2% BSA. 400 ng/ml CSP was added to the bacterial suspension (CSP-1 for ST5435, CSP-2 for ST700; Cambridge Biosciences). Suspensions were incubated at room temperature (RT) for 5 minutes, mixed with 300-500 ng transforming DNA, and further incubated at 37°C with 5% CO_2_ for 2 hours. The entire suspension was then plated on blood agar plates supplemented with relevant antibiotics. Transformants were screened using colony PCR and confirmed by Sanger sequencing. A complete list of plasmids and primers used in this study is listed in Tables 2 and 3.

**Table 3.**
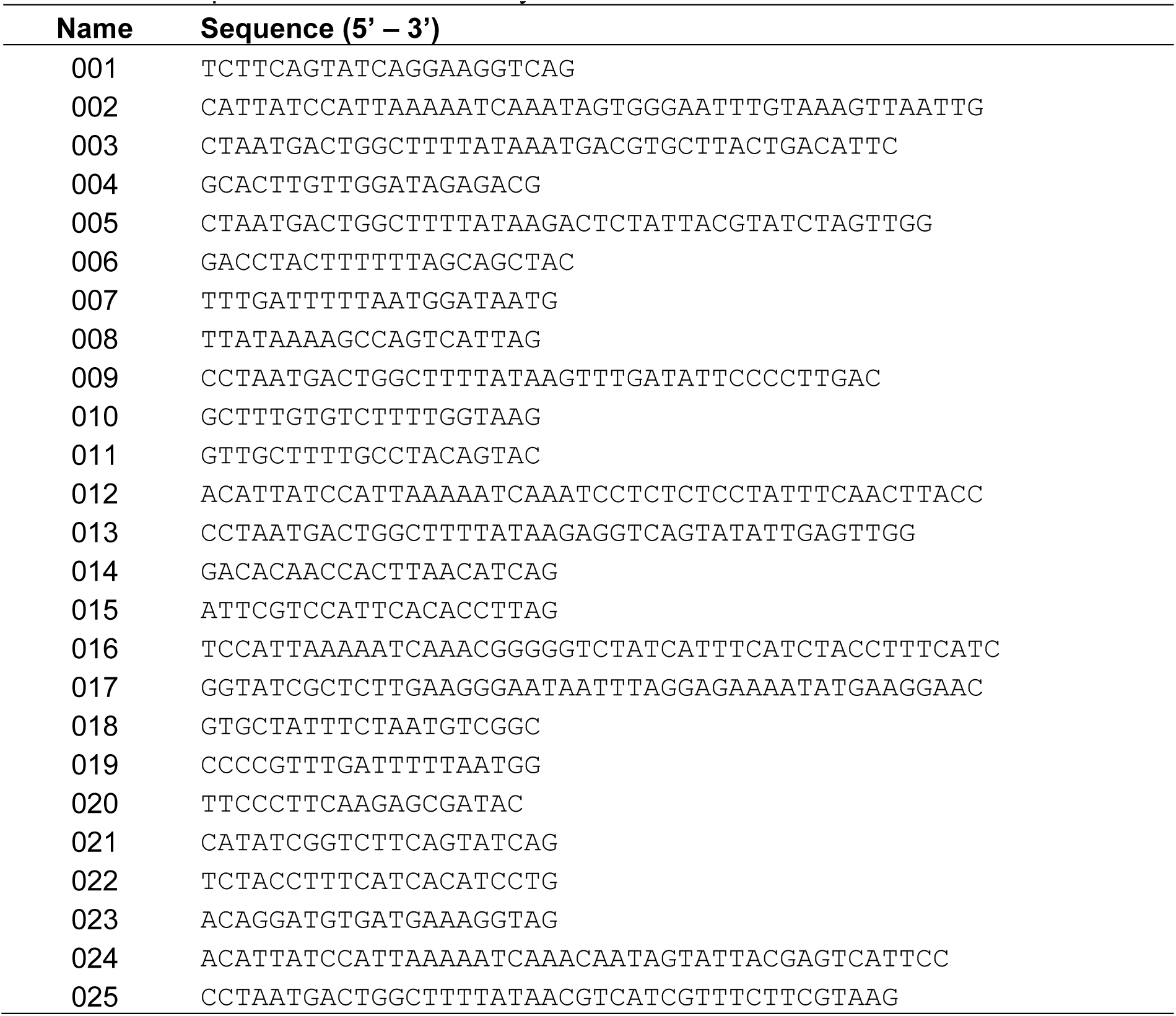
List of primers used in this study.

To construct the transforming DNA for making the unencapsulated (*cps::cat*) mutants, the areas flanking the capsular locus were amplified using primer pairs 001/002 and 003/004 from WT ST700 genomic DNA and primer pairs 001/002 and 005/006 from WT ST5435 genomic DNA. The chloramphenicol resistance cassette (*cat*) was amplified from pEMcat using primer pairs 007/008. PCR products corresponding to the upstream and downstream regions and *cat* were joined using overlap extension PCR (OE-PCR) with primer pairs 001/004 or 001/006, respectively. The purified OE-PCR product was used in transformation assays. Transforming DNA for making the *acl*-mutant (*acl::cat*) was constructed as described above using primer pairs 001/002 and 009/010 to amplify the up/downstream region of *acl* from ST5435 and primer pair 001/010 to generate the OE-PCR product. Transforming DNA for making the *cpsD* mutant (*cpsD::cat*) was constructed similarly using primer pairs 011/012 and 013/014 for the initial PCR and primer pair 011/014 for the OE-PCR product. Transforming DNA for making the *cpsB* mutant (*cpsB::spec*) utilized primer pairs 015/016 and 017/018 to amplify the flanking region, and primer pair 015/018 for the OE-PCR. Instead of *cat,* a spectinomycin resistance cassette (*spec*) was amplified from pR412 using primer pair 019/020 and used in OE-PCR.

To reintroduce *acl* into ST700 (*acl+*), the entire *acl* region was amplified in 3kb fragments from WT ST5435 genomic DNA using primer pairs 021/022 and 023/024. The downstream region, which encompasses part of the coding region for Ugd, was amplified from WT ST700 genomic DNA using primer pairs 025/010. The three PCR pieces and the *cat* cassette were assembled using NEB Gibson Assembly master mix to obtain the transforming DNA.

### Opsonophagocytic killing assay (OPK)

OPK analysis was performed using a WHO-approved standardized assay (37). Briefly, the strains were compared to assess the reduction in killing against a reference serotype 3 strain (an optochin-resistant variant of the ST180 strain Wu2) using pooled serum samples from adults vaccinated with PCV13. THY and 4% filtered fetal bovine serum were used for the assay. Strains were incubated with serial dilutions of pooled human serum prior to treatment with HL-60 cells, baby rabbit complement, and plating onto agar plates. The opsonic index was calculated as the reciprocal of the serum dilution reducing the bacterial colony forming units (CFU) to 50 of the no serum control.

### Complement deposition

Working stocks of *S. pneumoniae* were thawed and resuspended in 1X PBS to obtain ∼5 x 10^8^ CFU/ml. 10 µl bacterial suspensions (∼5 x 10^6^ CFU) were incubated with 10 µl 20% pooled human serum in 1x PBS for 30 mins at 37°C, washed 2x with 1x PBST (0.1% Tween-20) and labelled with 50 µl 1:300 FITC-polyclonal goat anti-human C3b antibody (ICN) on ice and in the dark for 30 mins. Bacterial suspension was washed 2x and fixed in 50 µl 3% PFA in PBS. Suspension volume was adjusted up to 250ul with 1x PBS prior to injection into the flow cytometer (BD (Beckton-Dickinson) FACSVerse).

### Growth curves

Overnight growth assays were performed using a BioTek Synergy H1 Plate Reader at 37°C, 5% CO_2_. Working stocks were thawed, harvested, and resuspended in BHI to a starting OD_600_ of 0.1, after which 200 µl were dispensed into a 96-well plate in technical triplicates. The plate was then incubated in the plate reader, and OD_600_ readings were taken every 15 minutes for a minimum of 18 hours. The growth curves presented are an average of three experiments conducted on three separate days.

### Collection of shed and attached pneumococcal capsule

Pneumococcal strains were grown in BHI at 37°C with 5% CO_2_ from a starting OD_600_ of 0.1 to an OD_600_ of 0.3. A 1 ml aliquot was harvested by centrifugation at 8,000 x *g* for 8 minutes, and the supernatant (“shed capsule”) and pellet (“attached capsule”) were separately stored at -70°C. The remainder of the culture was split into two and supplemented with either 20% normal human serum (NHS; Merck, S1-100ml) or an equivalent volume of BHI (no serum). The 20% NHS and no serum cultures were incubated at 37°C with 5% CO_2_ for an additional hour. After this, a 1 mL fraction was removed and subjected to harvesting as described above. 10 µl aliquots were taken at the start of the growth phase, at an OD_600_ of 0.3, and after an additional 1-hr incubation for CFU enumeration. These CFU counts were also used to evaluate bacterial survival in the presence of serum. The experiment was repeated at least three times on separate days.

### Quantification of shed and attached capsule

Quantification of the capsule was performed using a modified protocol from Chun *et al*. (62). For shed capsule samples, the thawed supernatant was centrifuged at 8,000 x *g* for 8 mins to ensure minimal carryover of intact cells. 250 µl of supernatant was transferred to a fresh microcentrifuge tube, incubated with 0.25 mg proteinase K (Qiagen) at 50°C for 1 hour and centrifuged at 13,000 x *g* for 2 mins to remove cellular debris. 100 µl aliquots of supernatant were transferred in duplicate to an ELISA plate and diluted 2-fold in 50 mM carbonate buffer, pH 9.6. Each well contains a final volume of 50 µl.

For the attached capsule samples, thawed pellets were resuspended in 1 mL of 1x PBS. After this, 250 µl was transferred to a fresh tube and treated with proteinase K as described above. The suspension is centrifuged at 13,000 x *g* for 2 minutes, and the pellet is resuspended in an equivalent volume of carbonate buffer. 100 µl aliquots were transferred in duplicate to an ELISA plate and diluted as described above.

Cross-adsorbed primary antibody was prepared by incubating Pneumococcus Type 3 serum (rabbit) at a 1:2,000 dilution in blocking buffer with 35 µl heat-killed ST700 *cps::cat* and 35 µl heat-killed ST5435 *cps::cat* suspensions per µl of serum used, overnight at 4°C statically. The heat-killed suspensions were prepared from cultures harvested at OD_600_ of 0.8, heated killed at 65°C for 45 minutes, and concentrated 5X. The antibody suspension was centrifuged at 3,000 x *g*, 5 minutes to remove heat-killed cells prior to use.

ELISA plates were coated with 2-fold dilutions of experimentally obtained capsule suspensions and purified serotype 3 polysaccharide capsule (Oxford Biosystems; standard range 10-640 ng/ml) at 4°C overnight, with rocking. Plates were then blocked with 1% BSA in PBS for 1 hour at RT with rocking, washed four times in 1x PBST (0.05% Tween-20), and incubated with adsorbed primary antibody for 1 hour at RT with rocking (Oxford Biosystems). The plate was then washed 4X, incubated with 1:10,000 secondary antibody (goat anti-rabbit IgG conjugated to HRP; LifeTech) in blocking buffer for one hour at RT with rocking, washed 4X, and incubated with SureBlue TMB Peroxidase substrate (Insight Biotech) until adequate color change was observed in the standard. An equivolume of 1N HCl was added to stop the reaction, and absorbance readings were taken at 450nm. The amount of polysaccharide capsule in each sample was calculated using the standard curve generated and normalized to CFU counts.

### Phosphoproteomics

Pneumococcal strains grown to OD_600_ of 0.3 in THY were treated with 20% normal human serum (Merck; 20% NHS) or THY (0% serum) for 1 hr at 37°C, 5% CO_2_, and then harvested via centrifugation at 3,000 x g for 15 mins. Cell pellets were resuspended in cold lysis buffer (0.1% NP-40 in 1X PBS (pH7.4), supplemented with 50 mM β-glycerol phosphate, 10 mM sodium orthovanadate and cOmplete Mini Protease Inhibitor Cocktail (Roche) and transferred to homogenizer tubes containing glass beads (Stratton Scientific). Pneumococci were lysed by bead-beating (6,200 RPM, 45 seconds on, 20 seconds off, 4 cycles), treated with benzonase for 1 hour on ice to remove nucleic acids, and centrifuged at 2,400 x *g* for 10 mins to remove cellular debris. Sample collection was repeated twice on different days, resulting in a total of 3 biological replicates. Protein lysates were saved at -70°C.

Thawed lysate samples were reduced with 10 mM dithiothreitol (DTT) for 1 hour at RT and then alkylated with 20 mM iodoacetamide for 30 minutes at RT. The alkylation reaction was quenched with an additional 20 mM DTT and diluted with 50 mM HEPES to reduce the urea concentration to <2M prior to digestion. Proteolytic digestion was carried out by the addition of 4 μg LysC (WAKO) and incubated at 37°C for 2.5 hrs followed by 10 μg trypsin (Pierce) treatment overnight at 37°C. After acidification, C18 MacroSpin columns (Nest Group) were used to clean up the digested peptide solutions and the eluted peptides dried by vacuum centrifugation. Samples were resuspended in 50 mM HEPES and labelled using the 0.8 mg Tandem Mass Tag 10plex isobaric reagent kit (Thermo Scientific) resuspended in acetonitrile. Labelling reactions were quenched with hydroxylamine and a pool was made of each set of samples. Acetonitrile content was removed from the pooled TMT solution by vacuum centrifugation. Labelled peptide pools were acidified and then cleaned using a Sep-Pak C18 (Waters). The eluted TMT-labelled peptides were dried by vacuum centrifugation, and phosphopeptide enrichment was subsequently carried out using the SMOAC (Sequential Metal Oxide Affinity Chromatography) strategy with High Select TiO2 and Fe-NTA enrichment kits (Thermo Scientific).

Eluates were combined prior to fractionation with the Pierce High pH Reversed-Phase Peptide Fractionation kit (Thermo Scientific). The dried TMT-labelled phosphopeptide fractions generated were resuspended in 0.1% TFA for LC-MS/MS analysis using a U3000 RSLCnano system (Thermo Scientific) interfaced with an Orbitrap Fusion Lumos (Thermo Scientific). Each peptide fraction was pre-concentrated on an Acclaim PepMap 100 trapping column before separation on a 50 cm, 75μm I.D. EASY-Spray Pepmap column over a 3-hr gradient run at 40°C eluted directly into the mass spectrometer. The instrument was run in data-dependent acquisition mode with the most abundant peptides selected for MS/MS fragmentation. Two replicate injections were made for each fraction with different fragmentation methods based on the MS2 HCD and MSA SPS MS3 strategies described in Jiang *et al*. (63).

The acquired raw mass spectrometric data were processed in MaxQuant (version 1.6.2.10) for peptide and protein identification. The database search was performed using the Andromeda search engine against predicted protein sequences from BVY11Z and BYV23H whole-genome sequences (64, 65). Fixed modifications were set as Carbamidomethyl (C) and variable modifications were set as Oxidation (M) and Phospho (STY). The estimated false discovery rate was set to 1% at the peptide, protein, and site levels. A maximum of two missed cleavages was allowed. Reporter ion MS2 or Reporter ion MS3 was appropriately selected for each raw file. Other parameters were used as preset by default in the software. The MaxQuant output file PhosphoSTY Sites.txt, an FDR-controlled site-based table compiled by MaxQuant from the relevant information about the identified peptides was imported into Perseus (v1.4.0.2) for data evaluation (66). Volcano plots were constructed using ggplot2 (67).

The mass spectrometry proteomics data have been deposited to the ProteomeXchange Consortium (http://proteomecentral.proteomexchange.org) via the PRIDE partner repository with the dataset identifier PXD054709 (68, 69).

### Ethical approval

PCV13-immunized serum samples were initially collected for an experimental human pneumococcal challenge trial approved by the UK National Health Service Research Ethics Committee (Ref: 12/NW/0873) (36). Leftover PCV-immunized serum samples were pooled and used in the OPK assay. Pooled serum samples used in flow cytometry experiments were collected with the ethical approval granted by the University College London (UCL) Institutional Review Board (3078/001). All donor information was anonymized to ensure confidentiality and privacy.

## ACKNOWLEDGEMENTS

We would like to thank DM Ferreira for the pooled immunized human serum and CJ Yang for expert support with ggplot2. This study was funded by a Medical Research Council grant (MR/T016329/1) awarded to RSH, a Wellcome Investigator award (reference 221803/Z/20/Z) to JSB, as well as a UCL Fellowship Incubator Award and a UCL Division of Infection and Immunity ECR Innovation Award awarded to JMC. Work was undertaken at UCLH/UCL which received funding from the Department of Health’s NIHR Biomedical Research Centre’s funding scheme. Phosphoproteomics work was supported by The Francis Crick Institute’s Proteomics facility, with special thanks to M Skehel. RSH is an NIHR Senior Investigator. The views expressed are those of the authors and not necessarily those of the NIHR.

